# BaCNet: Deep Learning Accelerates Novel Antibiotic Discovery Against Resistant Pathogens

**DOI:** 10.1101/2025.02.26.640477

**Authors:** Yuki Otani, Daisuke Koga, Yasunari Wakizaka, Hideyuki Shimizu

## Abstract

Drug-resistant infections pose a global health challenge and necessitate the rapid development of novel antibiotics. Although high-speed and high-accuracy *in silico* drug discovery methods using AI have been established, only a few approaches that specifically target antibiotic development have been developed. This gap significantly limits our ability to rapidly discover effective antibacterials against emerging resistant pathogens. Here, we have developed BaCNet, an AI system that accurately predicts the binding affinity between bacterial proteins and compounds using only amino acid sequences and compound SMILES representations. Our approach integrates a protein language model with three complementary compound embedding methods, achieving high prediction accuracy and effectively maintaining performance when tested on previously unseen bacterial species. BaCNet successfully rediscovered known antibiotics and identified promising novel candidates, with molecular dynamics simulations confirming stable binding of top hits. Moreover, by integrating a compound generation and optimization system with BaCNet, we discovered novel compounds not present in existing databases with significantly enhanced predicted antibacterial activity. BaCNet represents a promising platform that could accelerate the identification of urgently needed treatments against resistant pathogens.

## Introduction

Antimicrobial resistance (AMR) poses a serious threat to global public health, undermining the foundations of modern medicine^1–3^. In particular, the widespread dissemination of multidrug-resistant bacteria, including those encompassed by the ESKAPEE group (i.e., *Enterococcus faecium*, *Staphylococcus aureus*, *Klebsiella pneumoniae*, *Acinetobacter baumannii*, *Pseudomonas aeruginosa*, *Enterobacter* spp., and *Escherichia coli*), as identified by the World Health Organization, has significantly narrowed available therapeutic options, resulting in increased morbidity, mortality, and escalating healthcare costs^4,5^. Although the development of novel antimicrobial agents is urgently needed, traditional drug discovery processes are both time-consuming and costly, requiring an average of approximately 12 years and US$1.8 billion to bring a single new drug to the market^6,7^. Consequently, responding swiftly to emerging infectious threats has become increasingly challenging. Notably, two-thirds of the antimicrobial agents currently in use were developed between the 1940s and the 1960s, and in recent years, there has been little progress in the development of antimicrobials with novel mechanisms of action^8–10^.

Recent advances in computational science, particularly in artificial intelligence (AI), have provided innovative methods that accelerate the drug discovery process and offer the potential to address current challenges^11–13^, while also garnering significant attention from the pharmaceutical industry^14,15^. Current AI-driven drug discovery approaches include *de novo* drug design for generating novel compounds^16,17^, structure prediction methods exemplified by AlphaFold2^18^ and RoseTTAFold^19^, and techniques for predicting the binding affinities between target proteins and compounds^20–22^. However, these approaches were primarily developed to target human proteins. Bacterial proteins, on the other hand, possess transcriptional and post-translational modification systems that differ from those of eukaryotes, making it challenging to directly apply these methods to anti-infective drug development^23,24^. Consequently, a systems-level understanding of protein–compound interactions is imperative for the identification of bacterial drug targets and the development of effective therapies.

In recent years, rapid advances in large language models (LLMs) have spurred the development of protein language models (PLMs), which conceptualize protein amino acid sequences as a form of language, thereby driving transformative innovations in medicine and the life sciences^25^. Among these, Evolutionary Scale Modeling-2 (ESM-2)^26^ represents a large-scale PLM trained on protein sequences from a diverse array of organisms, including bacteria, and holds promise for capturing the unique characteristics of microbial proteins that conventional models have struggled to discern. Indeed, ESM-2 has demonstrated its efficacy in tasks such as classifying bacterial toxin proteins^27^ and predicting the evolutionary trajectory of the SARS-CoV-2 spike protein^28^. These advances underscore the potential of PLMs for modeling complex biological systems at the molecular level.

In response to the urgent need for novel antimicrobial agents, we have developed an AI-driven drug discovery system, BaCNet (**Ba**cterial **C**ompound-protein interaction **Net**work), specifically tailored for the identification of antimicrobial candidates (Fig. 1). BaCNet is a deep learning model trained on over 16 million compound–protein interactions (CPIs) derived from ESKAPEE pathogens. By leveraging ESM-2 as its foundation, BaCNet captures the structural and physicochemical properties of bacterial proteins, enabling a comprehensive and rapid exploration of vast chemical spaces for potential antimicrobial compounds (Fig. 1a). We demonstrated that BaCNet can successfully rediscover known antimicrobial agents and identify novel compounds with a high likelihood of exhibiting antimicrobial activity. Furthermore, molecular dynamics simulations confirmed the binding affinity of the identified novel compounds with their target proteins. Additionally, by integrating an *in silico* compound generation and optimization system with BaCNet, we have expanded the exploration space beyond existing compound libraries and successfully generated novel compounds predicted to exhibit superior antimicrobial activity (Fig. 1b). Thus, BaCNet represents a powerful platform for accelerating the development of next-generation therapeutics against drug-resistant infections and is expected to contribute to preparedness for future pandemics.

**Figure 1.**
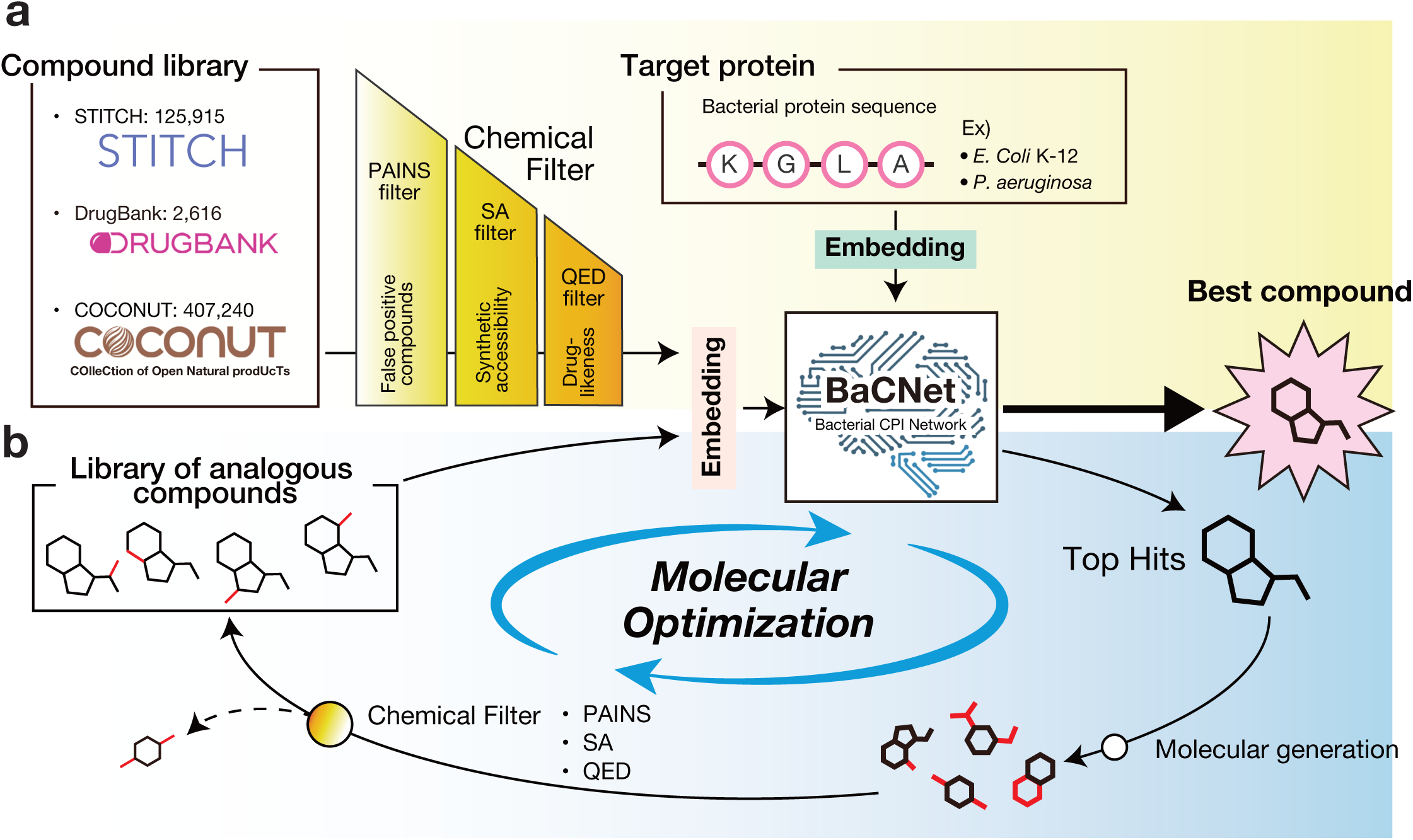
Overview of BaCNet for AI-driven antimicrobial drug discovery. **(a)** Screening Flow: BaCNet is a deep learning model designed to predict the binding affinity between bacterial proteins and compounds. In this study, BaCNet was used to screen compound libraries obtained from the existing databases (STITCH, DrugBank, and COCONUT) to identify potential antimicrobial candidates. The STITCH database provided compound–protein interactions (CPIs) data related to ESKAPEE pathogens, which were used as training data for BaCNet. Compound libraries were obtained from DrugBank (approved drugs) and COCONUT (natural products) and served as screening targets. **(b)** Optimization Flow: A compound generation and optimization system designed to enhance the predictive capacity of BaCNet. Starting with hit compounds as initial structures, an *in silico* compound generation model was employed to produce structurally similar analogs. The generated compounds were filtered using compound selection criteria (PAINS filter, SA filter, and QED filter) before undergoing re-evaluation by BaCNet for binding affinity. By iterating this cycle, novel compounds with high BaCNet scores can be generated efficiently.

## Results

### BaCNet learns binding affinity solely from bacterial protein sequences and compound SMILES representations

Although various methods for predicting protein-compound binding interactions have been proposed^29^, many of these approaches are based on training data from eukaryotic proteins, particularly human proteins, and repurposing these methods for anti-infectious drug discovery targeting prokaryotic proteins is challenging due to differences in post-translational modifications and other characteristics^23,24^. Therefore, we developed BaCNet (**Ba**cterial **C**ompound-protein Interaction **Net**work), an anti-infective drug discovery system trained on bacterial proteins. BaCNet takes only protein sequences and compound information, provided in SMILES (simplified molecular input line entry system) notation, as inputs and predicts the binding affinity between them, with its prediction score defined as the “BaCNet score”.

First, we collected the training data necessary to construct BaCNet from a public database (Fig. 2a). Specifically, we retrieved approximately 16 million protein-compound interaction (CPI) records from the large-scale database STITCH^30^, which compiles CPI information from a diverse array of organisms, including bacteria. We obtained data for 23 species belonging to the clinically important group of drug-resistant bacteria, ESKAPEE^4,5^. Subsequently, we partitioned the collected data. As an anti-infective drug discovery system, BaCNet must be capable of proposing effective antimicrobial agents for emerging infectious diseases. In other words, it must be able to predict the binding affinities of proteins from previously unseen bacterial species. To mimic the emergence of a novel infectious disease, we excluded the *E. coli* K-12 strain and *P. aeruginosa* from the dataset and designated them as external test data (Fig. 2a). Notably, because *P. aeruginosa* was not accompanied by other *Pseudomonas* species in the training data, it represents a taxonomically novel bacterial species at the genus level for BaCNet. These species were selected based on their clinical importance and the progression of antimicrobial resistance^31–34^. The remaining data were then divided into three sets: training (80%), validation (10%), and test (10%), with partitioning carefully designed to ensure that no protein sequences were duplicated across datasets, thereby mitigating the risk of overfitting.

**Figure 2.**
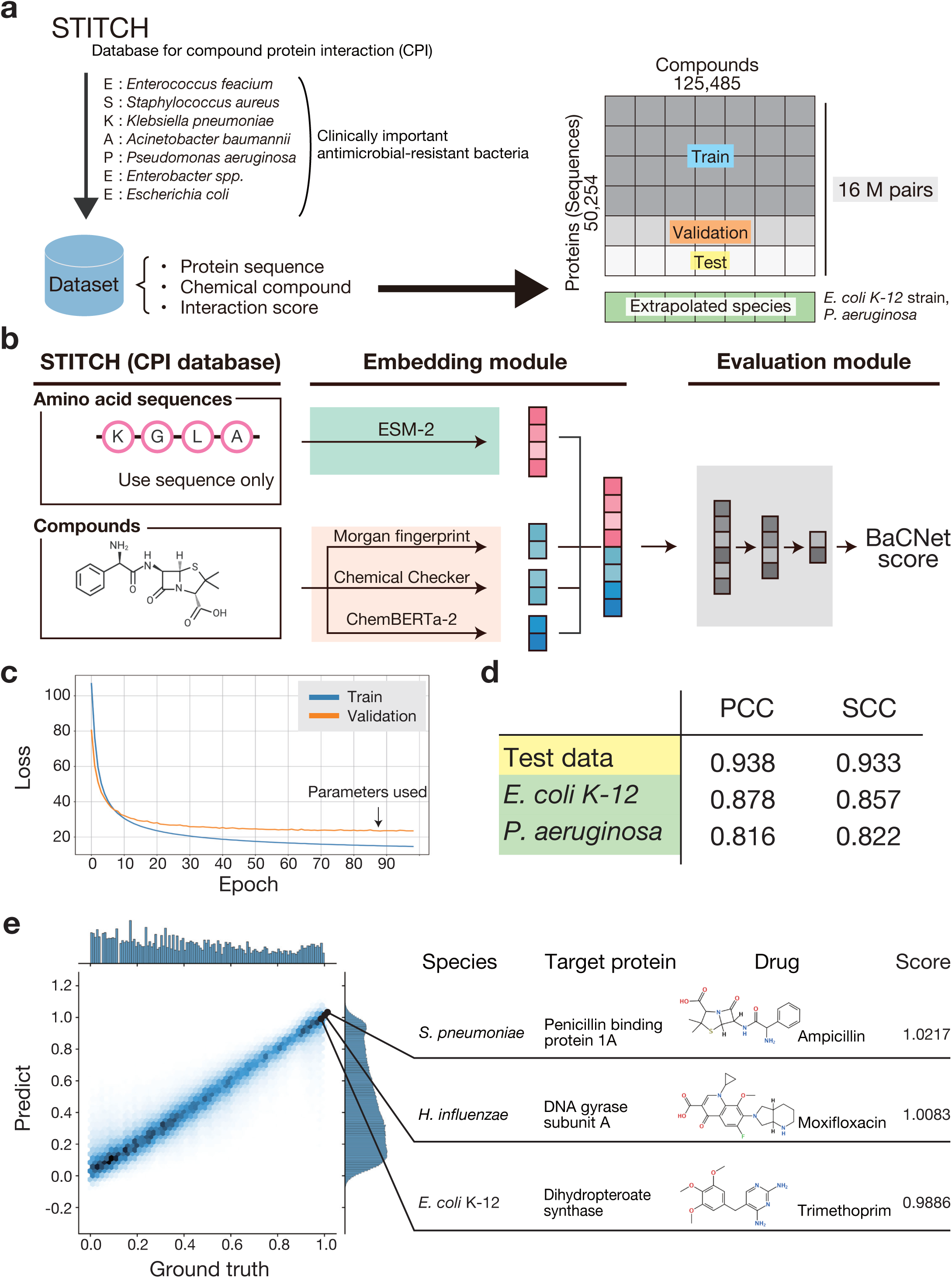
BaCNet accurately predicts the binding of bacterial proteins and compounds without 3D information. **(a)** Overview of the training dataset. Approximately 16 million CPIs data points derived from ESKAPEE pathogens were obtained from the STITCH database and were used for training. To simulate the emergence of a novel infectious disease, data corresponding to *E. coli* K-12 strain and *P. aeruginosa* were separated as an external test set. The remaining data were divided into training (80%), validation (10%), and test (10%) sets to ensure that no protein sequences were duplicated across these datasets. **(b)** BaCNet model architecture. BaCNet comprises two main modules: protein and compound embedding modules and an evaluation module. In the embedding module, protein sequences are embedded into a 5120-dimensional vector using ESM-2 (esm2_t48_15B_UR50D). Compounds are converted from SMILES notation into three separate vectors using Morgan fingerprint (1024 dimensions), Chemical Checker (10 signatures from A1–A5 and B1–B5, totaling 1280 dimensions), and ChemBERTa-2 (384 dimensions), which are then concatenated into a 7808-dimensional vector. In the evaluation module, the concatenated embedding vector is input into a multilayer perceptron (MLP) composed of three fully connected layers with 1024, 256, and 32 units. Following the application of the ReLU activation function and batch normalization, a single continuous output, defined as the BaCNet score (typically ranging from 0 to 1), is produced. **(c)** This panel displays the progression of the loss function (mean squared error, MSE) over epochs for both training (blue) and validation (orange) datasets. The horizontal axis indicates the number of epochs, and the vertical axis represents the MSE. Training was terminated when the validation loss did not improve for 10 consecutive epochs (indicated by an arrow). **(d)** The predictive performance of BaCNet was evaluated on both the test dataset and external test dataset (comprising *E. coli* K-12 strain and *P. aeruginosa*). The performance metrics included Pearson’s correlation coefficient (PCC) and Spearman’s correlation coefficient (SCC). **(e)** The left panel shows a scatter plot of BaCNet’s predicted values (vertical axis) against the true values (horizontal axis) for individual protein–compound pairs in the test dataset. The right panel shows the BaCNet scores for known pairs of antimicrobial agents and their target proteins (from *S. pneumoniae*, *H. influenzae*, and *E. coli* K-12 strain) obtained from DrugBank.

Figure 2b illustrates the model architecture of BaCNet. For protein feature extraction, we adopted the state-of-the-art protein language model (PLM) ESM-2^26^. ESM-2 is a PLM pre-trained on a large dataset of protein sequences from a diverse array of organisms, including bacteria, and is capable of generating numerical representations (embedding vectors) from amino acid sequences that capture high-level information, such as three-dimensional structure, function, and physicochemical properties. We hypothesized that these protein embedding vectors generated by ESM-2 would be effective for CPI prediction in the context of infectious diseases. On the compound side, the input is provided in SMILES notation, and we combined multiple embedding techniques. Since the structure and properties of compounds are critical factors in drug discovery, methods focusing on these aspects have been proposed^35–38^. Accordingly, for *in silico* drug discovery using BaCNet, we reasoned that numerical vectors reflecting various types of information, including the three-dimensional structure, substructure details, and physicochemical properties of compounds, would be useful for predicting binding affinity with proteins. Specifically, we employed three methods: Morgan fingerprint^39^ to capture local compound structures, Chemical Checker^40^ to represent compound similarity based on structural and interaction information, and ChemBERTa-2^41^, a compound language model capable of extracting global features and patterns of compounds. This multimodal compound representation is expected to allow for complementary utilization of the information provided by each embedding method and to achieve a higher predictive performance than any single embedding approach.

### BaCNet is a promising system for antimicrobial drug discovery

Based on the design described above, we evaluated the performance of BaCNet. First, by monitoring changes in the loss function during training, we confirmed that BaCNet achieved robust generalization performance without overfitting (Fig. 2c). The final model parameters were selected at the point where the validation loss ceased to improve for 10 consecutive epochs. Next, we quantitatively assessed the predictive performance of BaCNet by using a hold-out test dataset. BaCNet achieved a Pearson correlation coefficient (PCC) of 0.9382 (95% confidence interval (CI): 0.9378–0.9387) and a Spearman correlation coefficient (SCC) of 0.9325 (95% CI: 0.9321–0.9330) for the test set CPIs (Fig. 2d). Moreover, the mean squared error (MSE) was 0.00975, and the mean absolute error (MAE) was 0.0577, demonstrating that BaCNet can accurately predict the binding affinity between bacterial proteins and compounds.

Furthermore, we conducted a detailed investigation into the impact of combining the three compound embedding methods (Morgan fingerprint, Chemical Checker, and ChemBERTa-2) on the performance of BaCNet. The results revealed that integrating all three methods yielded superior performance compared with using any single method or any combination of the two methods (Table 1). This finding underscores the importance of incorporating multiple embedding techniques, each capturing different aspects of compound features, to enhance predictive accuracy.

**Table 1.** The ablation study of compound embedding methods.

|  | PCC | SCC | MSE |
| --- | --- | --- | --- |
| MF | 0.921 | 0.914 | 0.0126 |
| CBERTa | 0.899 | 0.894 | 0.0159 |
| CC | 0.911 | 0.904 | 0.0141 |
| MF + CBERTa | 0.936 | 0.930 | 0.0101 |
| MF + CC | 0.932 | 0.926 | 0.0107 |
| CBERTa + CC | 0.926 | 0.920 | 0.0115 |
| MF + CBERTa+ CC | 0.938 | 0.933 | 0.0098 |

MF: Morgan fingerprint, CC: Chemical Checker, CBERTa: ChemBERTa2, PCC: Pearson's correlation coefficient, SCC: Spearman's correlation coefficient
The highest value for each metrics (with the lowest value for MSE) was highlighted in red.

Next, we evaluated the extrapolative capability of BaCNet using external test dataset by assessing its predictive performance for proteins derived from bacterial species (*E. coli* K-12 strain and *P. aeruginosa*) that were never encountered during training. This external test dataset was prepared to simulate the emergence of a novel infectious disease (Fig. 2a). The results confirmed that BaCNet maintained a consistent level of predictive performance even for these previously unseen bacterial species (Fig. 2d). These findings strongly suggest that BaCNet can be applied to antimicrobial drug discovery against novel pathogens, such as those emerging during new infectious disease outbreaks.

To further demonstrate the utility of BaCNet, we evaluated whether it could predict known combinations of antimicrobial agents and their target proteins, that is, whether it could successfully rediscover established antimicrobials. Specifically, we applied BaCNet to score pairs of antimicrobial agents with well-characterized molecular mechanisms of action and their corresponding target proteins (Fig. 2e). In this experiment, we used proteins from bacterial species NOT included in BaCNet’s training data (*Streptococcus pneumoniae*, *Haemophilus influenzae*, and *E. coli* K-12 strain). BaCNet generated high scores for these known antimicrobial–target pairs (Table 2). These results corroborate that BaCNet can accurately predict compounds with inhibitory activity, even for bacterial species absent from the training dataset. Collectively, our findings support the conclusion that BaCNet is a promising system for antimicrobial drug discovery against bacterial infections.

**Table 2.** BaCNet score for known combinations of antimicrobial agents and their target proteins.

| Species | Target protein | Drug name | BaCNet score | Reference |
| --- | --- | --- | --- | --- |
| *E. coli* K12 | DNA gyrase subunit A | Ciprofloxacin | 1.0063 | Ref <sup>74</sup> |
| *E. coli* K12 | Dihydrofolate reductase | Trimethoprim | 0.9886 | Ref <sup>75,76</sup> |
| *E. coli* K12 | Dihydropteroate synthase | Sulfamethoxazole | 0.9500 | Ref <sup>77</sup> |
| *E. coli* K12 | Penicillin-binding protein 1A | Cefazolin | 0.8571 | Ref <sup>78</sup> |
| *S. pneumoniae* | Penicillin-binding protein 1A | Ampicillin | 1.0217 | Ref <sup>79</sup> |
| *S. pneumoniae* | Penicillin-binding protein 1B | Cephalexin | 0.8740 | Ref <sup>79</sup> |
| *H. influenzae* | DNA gyrase subunit A | Moxifloxacin | 1.0083 | Ref <sup>80,81</sup> |
| *H. influenzae* | DNA gyrase subunit A | Levofloxacin | 0.9981 | Ref <sup>82</sup> |

### Novel antimicrobial drug discovery using BaCNet

In the event of a novel infectious disease outbreak, it is imperative to identify candidate therapeutic compounds that are effective against drug-target proteins from previously unencountered bacterial species. To simulate this scenario, we employed a drug repositioning strategy using existing drugs and conducted a search for novel antimicrobial candidates targeting the penicillin-binding proteins (PBPs) of the *E. coli* K-12 strain, bacterial species that were never encountered during the training of BaCNet (Fig. 3a). PBPs play an essential role in bacterial proliferation and are the targeted by β-lactam antibiotics^42,43^. However, the emergence and dissemination of β-lactamases, enzymes that cleave the β-lactam ring essential for the activity of these antibiotics, have led to an increase in resistant bacterial populations^32,33,44,45^.

**Figure 3.**
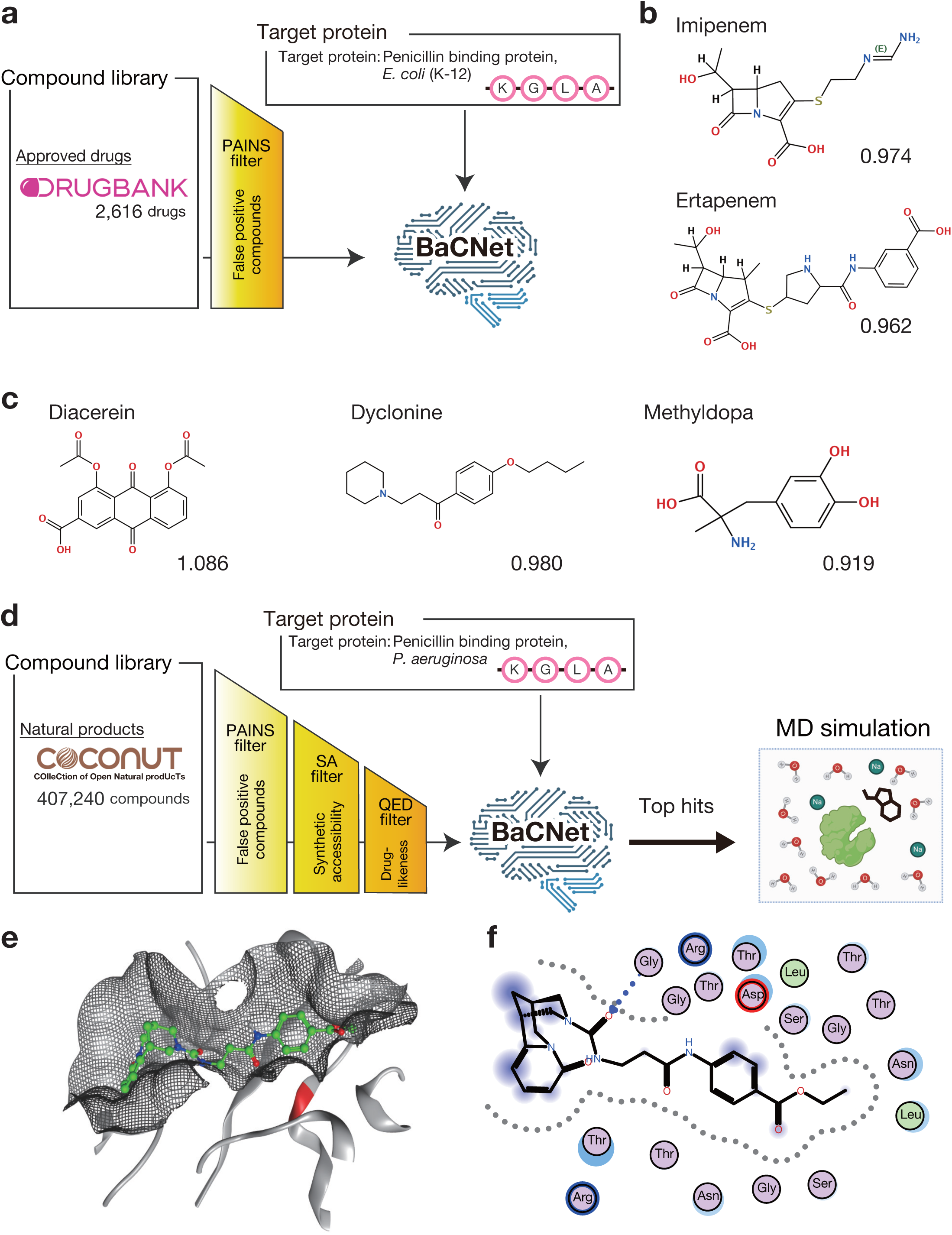
BaCNet identifies novel antimicrobial agents from established compound libraries. **(a)** Overview of the antimicrobial screening process using an approved drug library (DrugBank). Following the application of PAINS filter to remove false-positive compounds from the DrugBank database, screening was performed using BaCNet. **(b)** Structures and BaCNet scores of known antimicrobial agents identified among the top 30 compounds (ranked by BaCNet score) from DrugBank. **(c)** Structures and BaCNet scores of compounds previously reported to exhibit antimicrobial activity identified among the top 30 compounds (ranked by BaCNet score) from DrugBank. **(d)** Overview of the antimicrobial screening process using a natural compound library (COCONUT). Compounds were selected from the COCONUT database after applying PAINS filter, SA filter (excluding compounds with an SA score ≥ 5), and QED filter (excluding compounds with a QED < 0.4). These filtered compounds were then screened using BaCNet, and for the top 300 compounds (ranked by BaCNet score), binding stability with *P. aeruginosa* PBP was evaluated via molecular dynamics (MD) simulation. The SA filter assesses the synthetic accessibility, whereas the QED filter evaluates its suitability as an oral drug. **(e)** The complex structure of *P. aeruginosa* PBP (PDB ID: 4OON, shown in gray) with a top-ranked compound (COCONUT ID: CNP0419997, shown in green) identified from COCONUT screening. The active site residues of PBP are highlighted in red, and the mesh represents the contact surface between the protein and compound. **(f)** Two-dimensional depiction of protein–compound interactions within the complex shown in (e). Residues enclosed in the red box correspond to the active site of PBP, and the gray dashed line indicates the contact surface between the protein and compound.

Candidate compounds were selected from the DrugBank^46^ database after applying a Pan Assay Interference Compounds (PAINS) filter^47^ to exclude compounds prone to false positives due to their highly reactive functional groups. Scoring by BaCNet revealed that, among the top 30 compounds, in addition to known carbapenem antibiotics (imipenem and ertapenem, Fig. 3b) that target the PBP of the *E. coli* K-12, several drugs that have previously been suggested to possess antimicrobial activity (diacerein^48,49^, dyclonine^50^, and methyldopa^51^) were identified (Fig. 3c). Notably, the compound shown in Fig. 3c does not contain a β-lactam ring, suggesting that it may inhibit PBPs via a mechanism distinct from that of β-lactam antibiotics.

Next, we explored natural product-derived antimicrobial agents. Many antimicrobial agents developed thus far have been substances that organisms have acquired during evolution to defend against bacterial threats, suggesting that numerous undiscovered antimicrobial agents likely exist^52,53^. Accordingly, we screened for compounds expected to exhibit inhibitory activity against *P. aeruginosa* PBP using the natural compound library COCONUT^54^ (Fig. 3d). Prior to screening, we applied a PAINS filter, a Synthetic Accessibility (SA) score^55^ filter to evaluate the synthetic feasibility of the compounds, and a Quantitative Estimate of Drug-likeness (QED)^56^ filter to assess their suitability as oral drugs, thereby excluding compounds with undesirable properties (see Materials and Methods). A total of 186,036 compounds were obtained from COCONUT and screened for potential *P. aeruginosa* PBP inhibitory activity. Furthermore, we evaluated the validity of BaCNet’s predictions by performing molecular dynamics (MD) simulations on the top 300 compounds ranked by the BaCNet score. Specifically, we calculated the binding free energy of these compounds near the transpeptidase active site of the *P. aeruginosa* PBP and assessed protein-compound interactions to evaluate their potential as inhibitors (see Materials and Methods). Several compounds identified through BaCNet screening exhibited high binding stability to the *P. aeruginosa* PBP. Notably, one compound demonstrated a binding free energy of –9.14 kcal/mol and showed excellent complementarity with the shape of the PBP active site pocket (Fig. 3e, f). The stable binding of this compound to PBP^57,58^ suggests its potential as a novel non-β-lactam antimicrobial agent that does not rely on a β-lactam ring.

These lines of evidence show that BaCNet can rapidly and accurately identify candidate compounds for previously unencountered bacterial proteins, suggesting its potential as a powerful platform to support the rapid development of antimicrobial agents against emerging infectious diseases.

### Integrating compound generation and optimization systems with BaCNet enables the exploration of a vast chemical space

To rapidly provide effective therapeutics during a bacterial infection outbreak, it is essential to screen not only known compounds but also a vast number of potential candidate compounds. Hence, we developed an antimicrobial optimization system, termed “Optimization Flow”, which employs an *in silico* compound generation method to generate analogs of hit compounds and subsequently re-evaluate them using BaCNet (Fig. 4a). Optimization Flow expands the search space to include compounds not present in existing databases, thereby enabling a more comprehensive exploration. In Optimization Flow, new compounds are generated based on hit compounds; however, because some of these generated compounds may exhibit undesirable properties as therapeutic agents, we employed chemical filters (see Materials and Methods) to exclude them. The selected compounds were then re-evaluated using BaCNet.

**Figure 4.**
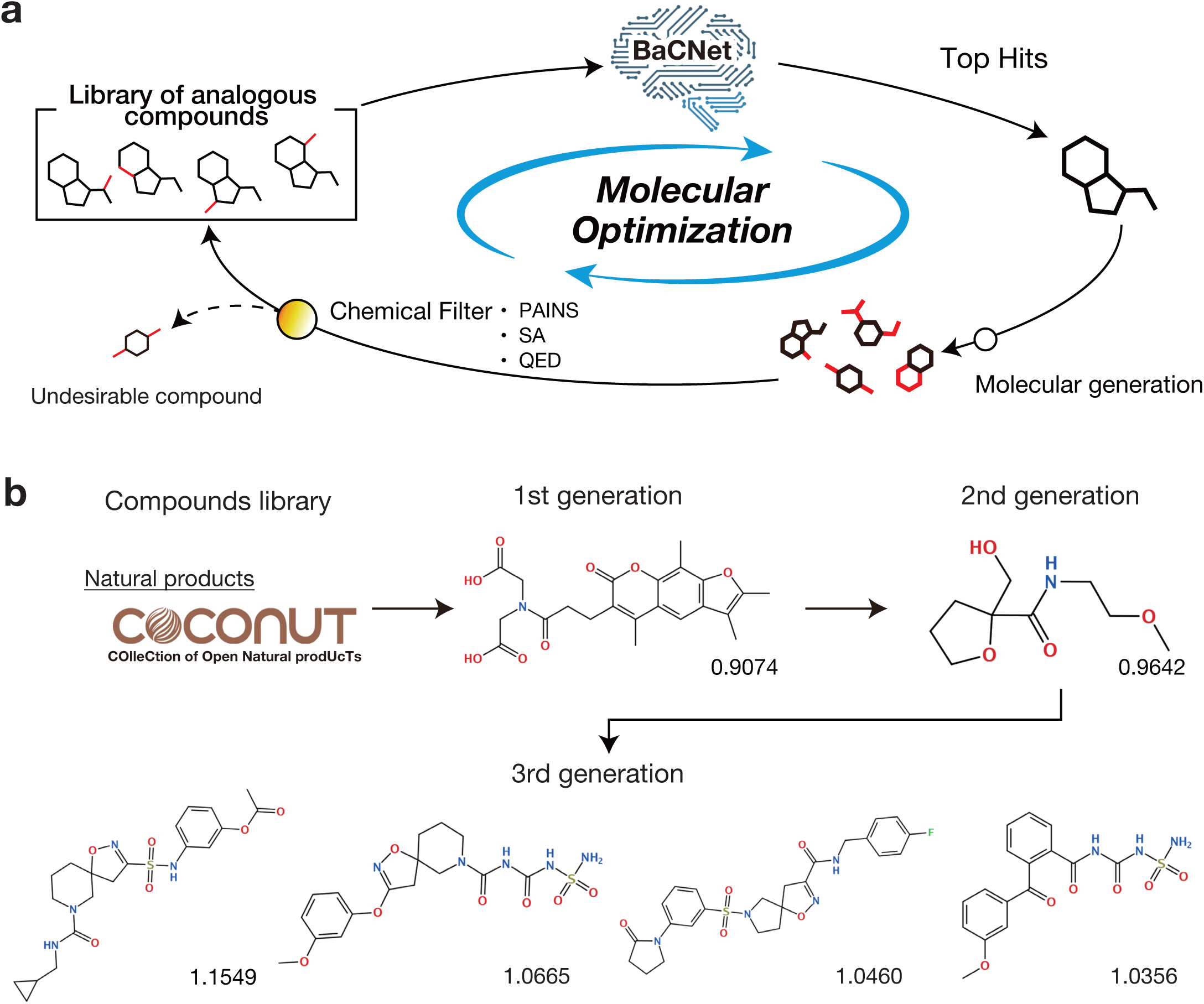
Incorporation of Optimization Flow into BaCNet enables the exploration of a vast chemical space. **(a)** Overview of the compound generation and optimization system (Optimization Flow). Hit compounds were used as initial structures, and a compound generation model is employed to produce structurally similar analogs. The generated compounds are then filtered using compound selection criteria (PAINS filter, SA filter, and QED filter) before being re-evaluated using BaCNet. By iterating this cycle, novel compounds with high BaCNet scores can be generated efficiently. **(b)** Starting with the hit compound (CNP0380562) obtained from the COCONUT database screening as the initial structure, three iterations of Optimization Flow were performed. The structures of the resulting compounds and their corresponding BaCNet scores are presented. Approximately 5,000 novel compounds were generated during each iteration.

We applied Optimization Flow to compounds initially screened from the COCONUT database. After repeating the cycle of generation and evaluation for three iterations, we successfully obtained multiple compounds with BaCNet scores significantly higher than those of the original compounds (Fig. 4b). Furthermore, among the compounds generated during the third iteration, the compound with the highest BaCNet score exhibited a score superior to that of any compound in the original COCONUT database. These results indicate that the introduction of Optimization Flow enables a comprehensive compound search independent of existing databases and enhances the quality of candidate antimicrobial agents.

## Discussion

Amid the escalating global threat posed by drug-resistant bacterial infections, the rapid development of novel antimicrobial agents has become an urgent challenge. Recently, *in silico* drug discovery leveraging artificial intelligence has emerged as a promising approach for addressing this issue. However, many existing AI-based drug discovery methods target human proteins and fail to consider the unique molecular mechanisms inherent to prokaryotic bacteria, thus limiting their applicability to antimicrobial development^23,24^. To overcome this limitation, we have developed BaCNet, a deep learning model that accurately predicts the interactions between bacterial proteins and compounds. BaCNet requires only sequence information as input, thereby providing an innovative, universal *in silico* platform for antimicrobial discovery that is independent of bacterial species.

The primary strength of BaCNet lies in its ability to predict the binding affinity between bacterial proteins and compounds with high accuracy and comprehensiveness. In validation experiments using external datasets not included in the training data, BaCNet maintained high accuracy for previously unseen bacterial species (i.e., *E. coli* K-12 strain and *P. aeruginosa*), demonstrating its potential as a universal platform capable of rapidly proposing antimicrobial candidates during emerging infectious disease outbreaks (Fig. 2d). Furthermore, in validation experiments employing known combinations of antimicrobial agents and their target proteins, BaCNet successfully rediscovered antimicrobial agents with diverse mechanisms of action with high accuracy (Fig. 2e, Table 2). Overall, these results indicate that BaCNet can accurately predict compounds with inhibitory activity, even for novel bacterial species, thereby supporting its reliability for practical antimicrobial drug discovery.

Using BaCNet for antimicrobial drug discovery, we successfully identified promising candidate compounds through both a drug repositioning strategy using existing drugs (DrugBank screening) and a natural compound strategy aimed at discovering novel antimicrobial agents (COCONUT screening) (Fig. 3). In the DrugBank screening, we identified not only known antimicrobial agents but also compounds previously suggested to possess antimicrobial activity (diacerein^48,49^, dyclonine^50^, and methyldopa^51^). The high BaCNet scores obtained for these compounds support the validity of our *in silico* antimicrobial screening approach. Furthermore, compounds identified via COCONUT screening were shown through MD simulations to bind stably to the active site of PBP (Fig. 3e, f). This binding mode differs from that of conventional β-lactam antibiotics and may render compounds less susceptible to degradation by β-lactamases. These findings suggest the potential for developing novel antimicrobial agents effective against β-lactamase–resistant bacteria, warranting further detailed investigation.

Furthermore, by developing and integrating a compound generation and optimization system, termed “Optimization Flow”, with BaCNet, we enabled the exploration of an expansive latent chemical space beyond that of existing compound libraries (Fig. 4). Utilizing Optimization Flow, we successfully obtained multiple novel compounds with significantly enhanced BaCNet scores compared to the original compounds (Fig. 4b), thereby demonstrating the efficacy of this approach for generating candidate antimicrobial agents.

However, this study has several limitations and presents avenues for future research. First, BaCNet does not explicitly incorporate three-dimensional structural information on proteins and compounds, meaning that its score may not directly reflect inhibitory activity. In addition to tertiary structural insights, considering the mode of action between proteins and compounds is crucial in drug discovery. With the advent of high-accuracy protein–compound complex structure prediction methods such as AlphaFold 3^59^ and RoseTTAFold All-Atom^60^, future studies integrating these structural data with functional insights into BaCNet could further enhance its predictive accuracy. Second, BaCNet does not account for critical drug properties such as pharmacokinetics and toxicity. Integrating BaCNet with *in silico* models that predict these properties^61,62^ may enable the development of a more comprehensive drug discovery support system. Future work should focus on enhancing BaCNet’s predictive performance, integrating safety evaluation systems, and experimentally validating predictions, thereby advancing BaCNet into a practical platform for antimicrobial drug discovery.

In conclusion, BaCNet, developed in this study, represents a novel and powerful *in silico* approach for accurately predicting the interactions between bacterial proteins and compounds, and is the first method specifically designed for this purpose. It facilitates high-accuracy screening of antimicrobial agents against previously unencountered pathogens and has the potential to serve as a robust framework in the fight against drug-resistant bacterial infections. We anticipate that this work will accelerate efforts to address the challenge of drug-resistant bacteria and strengthen preparedness against future pandemics.

## Materials and Methods

### Data acquisition for model training

All data used for training BaCNet were downloaded from Search Tool for Interacting Chemicals (STITCH, version 5.0) database^30^ (downloaded on May 31, 2023). In this study, from the information on 2,031 organisms available in the STITCH database, we extracted CPIs for 23 bacterial species associated with ESKAPEE and used these for model training. The protein sequences used for BaCNet training were unique sequences (i.e., no duplicates were allowed). Each CPI in the STITCH data was assigned a combined score based on experimental data, protein homology, evolutionary background, and literature evidence^30^. The combined scores were values with maximum value of 1000. Notably, the distribution of combined scores for protein-compound pairs is highly skewed toward values close to 0 (see Supplementary Fig. 1). To facilitate the learning process of BaCNet, these scores were transformed. First, to correct for the extreme skewness of the distribution, a Box-Cox transformation, which was designed to yield a distribution closer to the normal distribution, was applied. Then, the transformed scores were normalized to a range of 0 to 1.

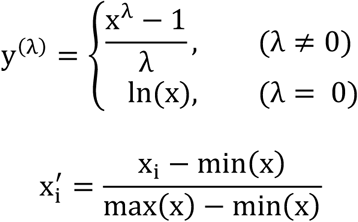

Here, the parameter *λ* was set to –1.4944. The Box–Cox transformation was applied using the boxcox function from the Python scipy.stats module (version 1.13.0). Under the assumption that the transformed data are normally distributed, the optimal *λ* parameter was determined by maximizing the corresponding log-likelihood function.

### Protein numerical representation using ESM-2

For the CPI data obtained from the STITCH database, protein sequences were converted into numerical representations that reflect their physicochemical properties using the ESM-2 model, specifically the esm2_t48_15B_UR50D configuration^26^. For each protein, matrix m_protein_ ∈ ℝ^length^ ^of^ ^sequence×5120^ was generated, where each amino acid was represented by a 5120-dimensional vector. A column-wise average was then computed to obtain *v̅*_*protein*_ ∈ ℝ^1×5120^, which was used as the protein feature vector for BaCNet.

### Compound numerical representation

Compounds obtained from STITCH were converted into numerical representations using SMILES strings, which preserve structural and characteristic information. The SMILES-transformed compound data were subsequently converted into numerical vector representations using three methods: Morgan fingerprint (MF)^39^, Chemical Checker (CC)^40^, and ChemBERTa-2 (CBERTa)^41^. These methods yielded vectors *v*_MF_ ∈ ℝ^1×1024^, *v*_*CC*_ ∈ ℝ^1×3200^, and *v*_CBERTa_ ∈ ℝ^1×384^, respectively. Using RDKit (version 2024.03.5)^63^, the Morgan fingerprint with a radius of 2 was computed from the compound SMILES strings, resulting in a 1024-dimensional binary vector. Chemical Checker^40^ provides an integrated feature representation that encompasses multiple levels of information, including compound structure, bioactivity, and target interactions, organized into 25 signatures (A1–A5, B1–B5, C1–C5, D1–D5, and E1–E5), each represented as a 128-dimensional vector. In this study, we utilized 10 of these signatures (A1–A5 and B1–B5), which reflect structural features and physicochemical properties, resulting in a combined 1280-dimensional vector for each compound. Morgan fingerprints encode the presence or absence of substructures in a binary format and have been widely used in drug discovery^64–66^. ChemBERTa-2^41^ applies LLM techniques to generate numerical vector representations that capture both the structure and activity of molecules. In this study, the Hugging Face Transformers library (version 4.33.3) was used to obtain a 384-dimensional embedding vector from ChemBERTa-2 (pretrained model: “DeepChem/ChemBERTa-77M-MLM”). Finally, the three embedding vectors were concatenated to form a single compound representation *v*_*compound*_ with a total dimensionality of 1024 + 1280 + 384 = 2688.

### Architecture and training of BaCNet model

BaCNet is a deep learning model that accepts amino acid sequences and SMILES strings as inputs, internally converting them into protein embedding representations via ESM-2 and compound embedding vectors, respectively, and then predicts the binding affinity score (termed the BaCNet score) between them (Fig. 2b). The input to BaCNet is a vector *v*_*input*_ ∈ ℝ^1×7808^ formed by concatenating these embedding vectors. The classification component of BaCNet consists of a multilayer perceptron (MLP) with three fully connected layers. At each layer, a weighted linear transformation was applied, followed by a rectified linear unit (ReLU) activation function and batch normalization. The number of units in the MLP layers was [1024, 256, 32], and the network produced a single output value, which was defined as the BaCNet score. Typically, the BaCNet score ranges from 0 to 1, with higher scores suggesting stronger binding affinity between the protein and compound. The mean squared error (MSE) was used as the loss function during training, defined as:

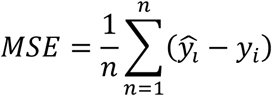

where *ŷ*_*i*_ and *y*_*i*_ represent the predicted and true CPI values, respectively, for the *i*th protein–compound pair, and training was conducted with a batch size of 32. The AdamW optimizer^67^ (weight decay = 0.001) was employed, and the learning rate was set to 0.0001. To mitigate overfitting, early stopping was applied if the loss in the validation dataset did not improve for 10 consecutive epochs. Model training and evaluation were implemented using PyTorch (version 2.1.0). Finally, the trained model was evaluated on a test dataset that was set aside prior to training, ensuring that it was only encountered during the final evaluation. The 95% confidence intervals (CIs) for the correlation coefficients were computed using 5,000 bootstrap samples. BaCNet was trained in parallel on eight NVIDIA A100 GPUs.

### Metrics for assessing model performance

To evaluate the predictive performance of BaCNet, we employed three metrics: Pearson’s correlation coefficient (PCC), Spearman’s correlation coefficient (SCC), and the coefficient of determination (*R*^2^). These metrics were defined by the following equations:

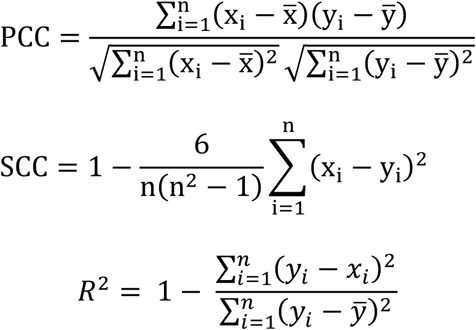

where *x*_*i*_ and *y*_*i*_ denote the predicted and true values, respectively, for the *i*th data point, while *x̅* and *y̅* represent the means of predvicted and true values, defined as 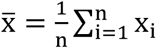 and 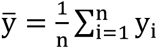 where n is the number of samples.

### Approved drugs and their target proteins

The drug–target protein pairs presented in Fig. 2e and Table 2 were collected from DrugBank^46^ (downloaded on June 24, 2024) based on documented reports. The SMILES representations of the compounds were retrieved from DrugBank, and the amino acid sequences of the target proteins were obtained from UniProt.

### *In silico* screening of novel antibiotics using BaCNet

From the approved drug database DrugBank^46^ (downloaded on June 24, 2024) and the natural compound library COCONUT^54^ (downloaded on May 14, 2024), we obtained 2,616 and 407,240 compound SMILES representations, respectively, which were used to screen bacterial protein inhibitors. Prior to screening, compound filters were applied to exclude compounds with undesirable properties as potential drugs. Specifically, the following filters were employed:

1. PAINS Filter: Compounds identified as Pan Assay Interference Compounds (PAINS)^47^ were removed using the rdkit.Chem.FilterCatalog module from RDKit (version 2024.03.5)^63^.
2. SA Filter: The Synthetic Accessibility (SA) score^55^ was calculated using RDKit’s rdkit.Contrib.SA_Scoremodule, and compounds with SA scores exceeding five were excluded.
3. QED Filter: The Quantitative Estimate of Drug-likeness (QED)^56^ was computed using RDKit’s rdkit.Chem.QED.qed function, and, following previous studies^68^, compounds with QED values below 0.4 were eliminated.

Only the compounds that passed these filters were screened using BaCNet.

### Molecular dynamics (MD) simulation and preprocessing

MD simulations were performed to evaluate the binding stability of the top-ranked compounds identified through BaCNet screening with the target protein. The Molecular Operating Environment (MOE, Chemical Computing Group Inc.) was used for MD simulations. Initially, the compounds, originally represented in SMILES format, were converted to SDF (structure-data file) format containing atomic coordinate information using the open-source RDKit library. Subsequently, MOE was employed to generate two-dimensional structures, add hydrogen atoms, and perform structural optimization using the Amber10:EHT force field^69^. The optimized structures were then utilized for both docking calculations and MD simulations.

The three-dimensional structure of *P. aeruginosa* PBP was obtained from the Protein Data Bank (PDB ID: 4OON, downloaded on October 7, 2024). MOE was used to add hydrogen atoms to this structure and to optimize it using the Amber10:EHT force field^69^. During preprocessing, extraneous small molecules were removed. Similar hydrogen addition and structural optimization were performed on the protein, which was then used for docking calculations. The docking site on PBP was identified using MOE’s SiteFinder, a tool that comprehensively detects potential docking sites based on protein structure and surface properties. The site used for docking between PBP and the compounds was selected based on UniProt annotation, ensuring its proximity to the active site.

### Docking calculations

Docking calculations were performed using MOE. Initially, the stability of the compound on the protein surface was evaluated using simulation with a time step of 1 fs. Once the energy had converged, the binding free energy between the protein and the compound was calculated. Thirty initial poses of the compound were generated using the Triangle Matcher placement algorithm^70^, and the calculations were repeated for each pose. The binding free energy was estimated using the London ΔG scoring function^71^, as shown in the following equation:

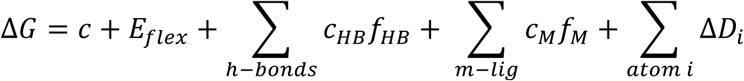

Where *c* represents the average loss of rotational and translational entropy of the compound, *E*_*flex*_ is the energy loss due to ligand flexibility calculated solely from its conformation, *f*_*HB*_ is a variable (ranging from 0 to 1) representing the geometric imperfection of hydrogen bonds, *C*_*HB*_ is the ideal hydrogen bond energy, *f*_*M*_ is a variable (ranging from 0 to 1) representing the geometric imperfection of metal-ligand interactions, *c*_*M*_ is the ideal energy for metal-ligand interactions, and *D*_*i*_ represents the desolvation energy of the *i*th atom of the ligand, which is calculated as follows:

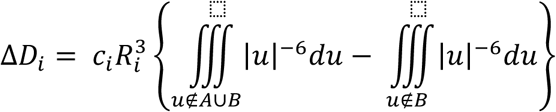

In this equation, *A* and *B* denote the spatial regions occupied by the protein and ligand, respectively, *R*_*i*_ is the solvation radius of the *i*th atom, and *c*_*i*_ is a constant specific to the atom type.

Additionally, the binding free energy was calculated using the GBVI/WSA ΔG function^72^, according to the following expression:

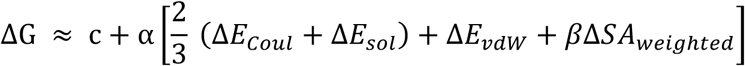

where *c* represents the average loss of rotational and translational entropy of the compound, *E*_*Coul*_ is the electrostatic energy, *E*_*sol*_ is the solvation electrostatic energy, *E*_*vdW*_ is the van der Waals energy, *SA_weig*hted*_* is the weighted solvent-accessible surface area, and *⍺* and *β* are constants.

### Compound generation and screening via Optimization Flow

To screen a broader range of candidate therapeutic compounds, we developed an integrated compound generation and optimization system termed Optimization Flow. Using NP-VAE^73^, a method that generates similar compounds from SMILES representations, 5,000 compounds were generated. Prior to screening, compound filters (PAINS filter, SA filter, and QED filter) were applied to exclude compounds with undesirable drug-like properties. Only the compounds that passed through these filters were screened using BaCNet.

## ACKNOWLEDGEMENTS

This work was supported by JST, PRESTO Grant Number JPMJPR22R6, Japan to H.S. This work was also partly supported by KAKENHI grants from the Japan Society for the Promotion of Science (JSPS) to H.S. (24H01755 and 23K18502), as well as Takeda Science Foundation. We thank H. Aso, S. Ohno, and other laboratory members for discussion and K. Tanaka for help with preparation of the manuscript.

## AUTHOR CONTRIBUTIONS

H. S. conceived of, designed, and supervised the study. Y.O. developed BacNet and performed all computational analyses, with inputs from D.K. and Y.W. Y.O. and H.S. jointly wrote the manuscript with the comments from all authors.

## Supplementary Information

**Supplementary Figure 1.**
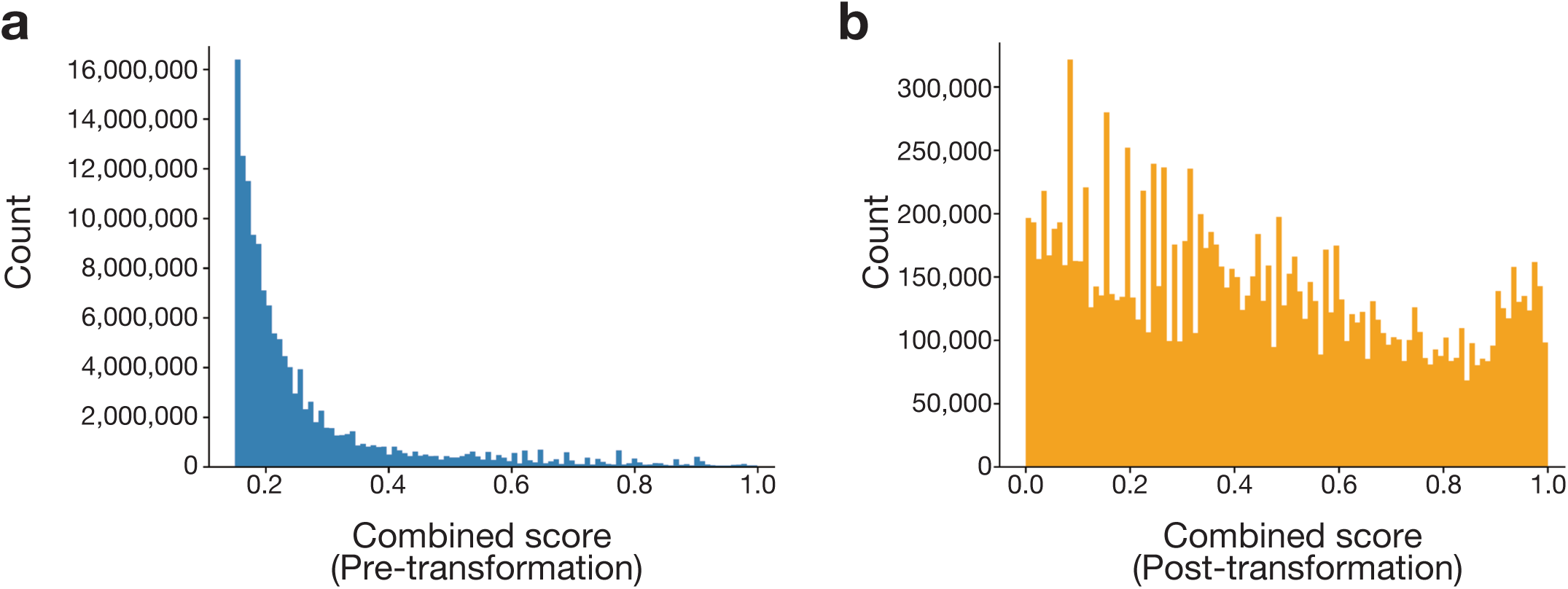
Distribution of the combined score. **(a)** Distribution of combined scores in raw training data. The x-axis represents the combined score, and the y-axis represents the frequency. (**b**) Distribution of combined scores in the training data after Box-Cox transformation and normalization. The x-axis represents the combined score, and the y-axis represents the frequency.

